# Comparative transcriptome and variant analyses of the pancreatic islets of a rat model of obese type 2 diabetes identifies a frequently distributed nonsense mutation in the lipocalin 2 gene

**DOI:** 10.1101/2024.09.13.609843

**Authors:** Norihide Yokoi, Naoki Adachi, Tomoki Hirokoji, Kenta Nakano, Minako Yoshihara, Saki Shigenaka, Ryuya Urakawa, Yukio Taniguchi, Yusaku Yoshida, Shigeo Yokose, Mikita Suyama, Tadashi Okamura

## Abstract

We have recently established the Zucker fatty diabetes mellitus (ZFDM) rat as a novel model of obese type 2 diabetes (T2D), originating from the obese Zucker fatty (ZF) rat harboring a missense mutation in the leptin receptor gene. Pathogenesis of dysfunction of the pancreatic islets and genetic factors of T2D in ZFDM rats remain unknown. Here, we perform comparative transcriptome and variant analyses of the pancreatic islets between the two strains. Among differentially expressed genes irrespective of obesity and glucose intolerance states, we identify a nonsense mutation, c.409C>T (p.Gln137X), in the lipocalin 2 (*Lcn2*) gene which encodes a secreted protein called neutrophil gelatinase-associated lipocalin, a well-known biomarker for inflammation. Interestingly, we find that the *Lcn2* mutation is distributed widely in rat species, such as commonly used DA and F344 strains. We examine the *Lcn2* mutation as a strong candidate gene for T2D in ZFDM rats by using genome editing of ZFDM rats in which the nonsense mutation is replaced with a wild-type nucleotide. We find that the genome editing well works but also observe that there is no significant difference in the development of T2D between genome-edited and original ZFDM rats. Finally, we perform a genetic linkage analysis by using backcross progeny between ZF and ZFDM rats and confirm that the *Lcn2* mutation exhibits no significant association with the onset of T2D. Our data indicate that several rat strains would serve as *Lcn2* deficient models, contributing to elucidate pathophysiological roles of Lcn2 in a wide variety of phenotypes.

## Introduction

Type 2 diabetes (T2D) is a multifactorial disease caused by insulin resistance and impaired insulin secretion from pancreatic β-cells; β-cell failure is a central element in the development and progression of type 2 diabetes. The β-cell failure in type 2 diabetes is a complex process involving multiple factors, such as insulin resistance and β-cell overload, lipotoxicity, chronic inflammation, oxidative stress, endoplasmic reticulum (ER) stress, and genetic and epigenetic factors. However, the precise mechanisms remain to be elucidated.

Transcriptome analysis of the pancreatic islets offers a powerful technique not only to reveal pathophysiological state of β-cells but to identify candidate genes in T2D. There have been several studies to identify candidate genes for T2D by transcriptome analyses in pancreatic islets from individuals with T2D and nondiabetic controls. Bacos et al. identified PAX5 as a potential transcriptional regulator of many T2D-associated differentially expressed genes (DEGs) in human islets (Bacos et al. 2023). Transcriptome profiling in db/db mice revealed that a reduction in the expression of Glut2, Ins1, Ins2, MafA, Mt1, and Pdx1 was indicative of dedifferentiation in db/db islets (Neelankal John et al., 2018). Griffen et al. (2001) demonstrated that the ZDF rat carries a genetic defect in beta-cell gene transcription, in which insulin promoter activity and insulin gene expression were reduced in even in lean animals. They concluded that the genetic reduction in beta-cell gene transcription in ZDF rats likely contributes to the development of diabetes in the setting of insulin resistance. Transcriptome profiling of ZDF rats showed an increase in the genes encoding proteases and extracellular matrix components that are associated with tissue remodeling and fibrosis (Zhou et al. 2005). In addition, ZDF rats showed an increase in vascular endothelial growth factor (VEGF)-A and Thrombospondin-1 genes, suggesting that an inability of the islet to maintain vascular integrity may contribute to beta-cell failure (Li et al. 2006). However, the primary genetic factors have not been clarified yet.

Among animal models of T2D, the Zucker fatty diabetes mellitus (ZFDM) rat has been derived from the obese Zucker fatty (ZF) rat harboring a missense mutation (*fatty, fa*) in the leptin receptor (*Lepr*) gene (Yokoi et al. 2013). Animals homozygous for the *fatty* mutation in both ZF and ZFDM strain exhibit obesity, whereas only male ZFDM rats develop T2D accompanied with histopathological changes in the pancreatic islets such as loss of islet structure and β-cell destruction (Yokoi et al. 2013; Gheni et al. 2015). In spite of the same origin, there is significant difference in genetic profiles between the two strains (Nakanishi et al. 2017); genetic factors involved in the development of T2D remain unknown.

We have previously performed transcriptome analysis of the pancreatic islets in ZFDM rats to examine the mechanism underlying functional differences between non-large and enlarged islets. Together with the insulin secretion experiment and metabolome analysis, we found that enlarged pancreatic islets show tumor cell-like metabolic features of glucose metabolism accompanied with reduced beta-cell specific gene expressions and glutamate production, which could contribute to the development of incretin unresponsiveness in obese T2D (Hayami et al. 2020). However, pathogenesis of dysfunction of the pancreatic islets and genetic factors involved in the development of T2D still need to be clarified.

In the present study, to elucidate gene expression profile of the pancreatic islets and primary genetic factors of T2D in the ZFDM rat, we performed comparative transcriptome analysis and variant search on the pancreatic islets between ZF and ZFDM rats. Among DEGs, we identified a nonsense mutation in a strong candidate gene for T2D, which is found to be existed frequently in rat species. We also evaluated the relevance of the mutation with T2D by both genome editing and genetic linkage analysis.

## Results

### Comparative transcriptome analysis of the pancreatic islets between ZF and ZFDM rats

To elucidate gene expression profile of the pancreatic islets and primary genetic factors of T2D in ZFDM rats, we performed comparative transcriptome analysis between ZF and ZFDM rats at 8 and 12 weeks of age (Fig. 1A). ZFDM *fa/fa* rats at 8 weeks of age showed normoglycemia with very slight glucose intolerance upon oral glucose load, while those at 12 weeks of age exhibited apparent glucose intolerance (Yokoi et al. 2013; Hayami et al. 2020). Since there was a large variation in size of the islets in *fa/fa* rats at 12 weeks of age (Hayami et al. 2020), we compared gene expression in non-large and enlarged (exceeding 300 μm in diameter) islets separately. Among ∼13,000 genes detected, we found DEGs for each comparison (Fig. 1B): 1,046 DEGs for ZF +/+ vs. ZFDM *fa/+* at 8 weeks of age (Supplemental Table S1), 359 DEGs for ZF *fa/fa* vs. ZFDM *fa/fa* at 8 weeks of age (Supplemental Table S2), 1,355 DEGs for ZF +/+ vs. ZFDM *fa/+* at 12 weeks of age (Supplemental Table S3), 778 DEGs for non-large islets of ZF *fa/fa* vs. ZFDM *fa/fa* at 12 weeks of age (Supplemental Table S4), 766 DEGs for enlarged islets of ZF *fa/fa* vs. ZFDM *fa/fa* at 12 weeks of age (Supplemental Table S5). Common DEGs at each age may represent primary expression difference between strains irrespective of obesity and subsequent glucose intolerance conditions, which may be reflected by primary genetic difference. We performed gene enrichment analysis on these common DEGs. Common 127 DEGs (up: 58, down: 69) at 8 weeks of age (Fig. 1C; Supplemental Table S6) were enriched in the GO term associated with “response to stimulus” and “extracellular matrix” (Supplemental Table S7). Common 166 DEGs (up: 114, down: 52) at 12 weeks of age (Fig. 1C; Supplemental Table S8) were enriched in the GO term associated with “positive regulation of multicellular organismal process”, “response to stimulus”, and “extracellular matrix” (Supplemental Table S9). Due to relatively small number of genes, there was no significant GO term for common 46 DEGs (up: 18, down: 28) for all the comparisons (Fig. 1C; Supplemental Tables S10, 11). However, these 46 DEGs may serve as strong candidate genes for T2D in ZFDM rats.

**Figure 1.**
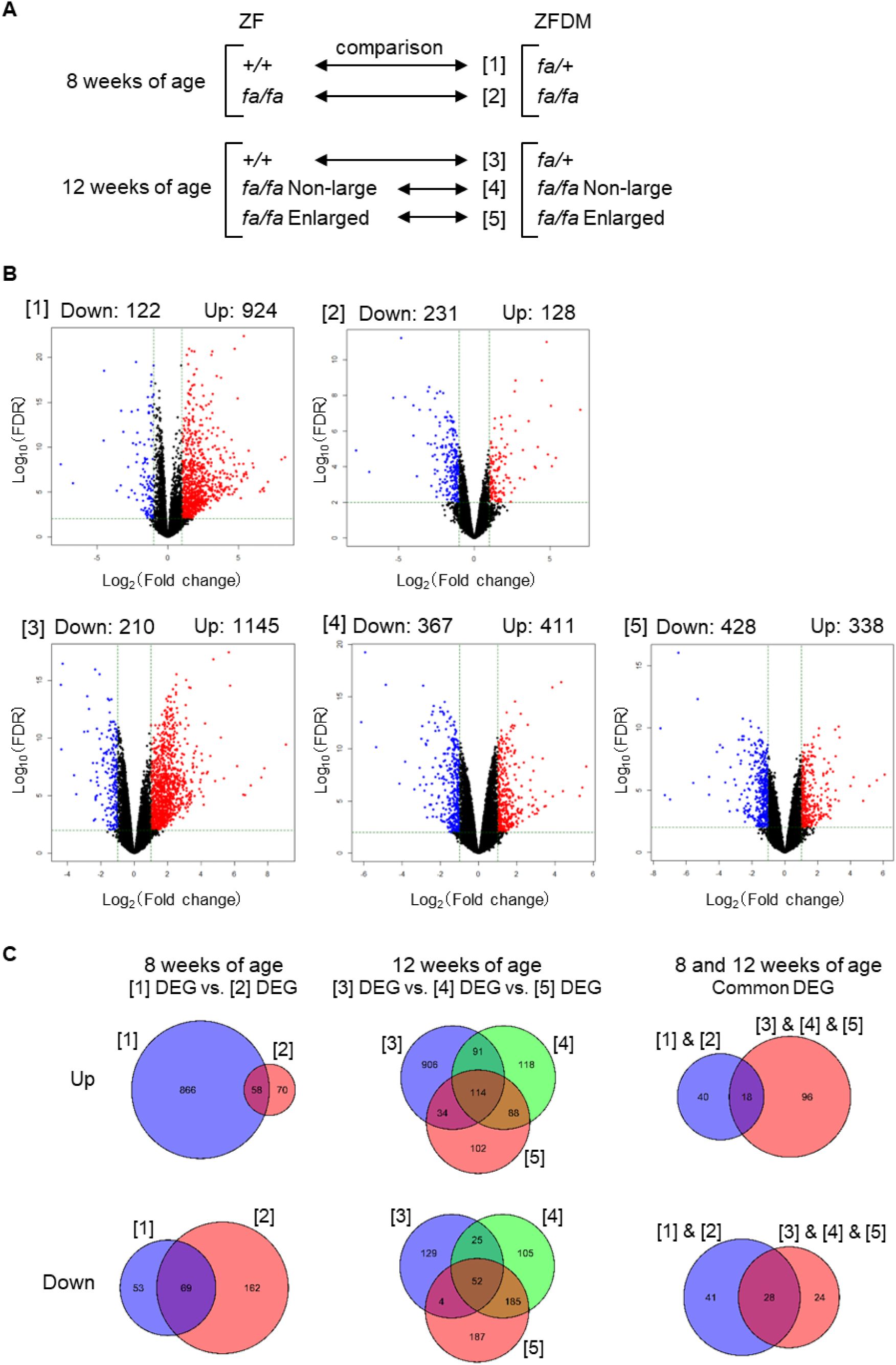
Comparative transcriptome analysis of the pancreatic islets between ZF and ZFDM rats. (*A*) Comparisons of transcriptome ([1] - [5]) between ZF and ZFDM rats at 8 and 12 weeks of age (n=3 each). (*B*) Volcano plots of the comparisons of transcriptome ([1] - [5]). Numbers of DEGs are shown as Down and Up in ZFDM rats as compared to ZF rats. (*C*) Venn diagrams showing common DEGs.

### Identification of a nonsense mutation in the *Lcn2* gene in the ZFDM rat

To further elucidate primary genetic factors of T2D in ZFDM rats, we performed variant search on RNA-seq data of the islets of ZF and ZFDM rats at 8 weeks of age and found that there were ∼5,900 variants including 5 stop-gained variants (3 genes) and 921 missense variants as compared with the rat reference genome sequence (Supplemental Table S12). The ZF rat has stop-gained variants in *Commd7* and *RT1-Db1* while the ZFDM rat has those in *Lcn2* and *RT1-Db1*. Among these genes, the lipocalin 2 (*Lcn2*) was included in the common 46 DEGs. In ZF rats, *Lcn2* gene expression was much higher in the islets of *fa/fa* rats as compared with that of +/+ rats, and the expression levels increased with age (Fig. 2A). In contrast, the expression was significantly lower in the islets of both *fa*/+ and *fa/fa* in the ZFDM rat as compared with those of ZF rats. As for ZF rats, the expression levels in ZFDM *fa/fa* rats were higher than those in ZFDM *fa*/+ rats, but the difference was much lower than that of ZF rats.

**Figure 2.**
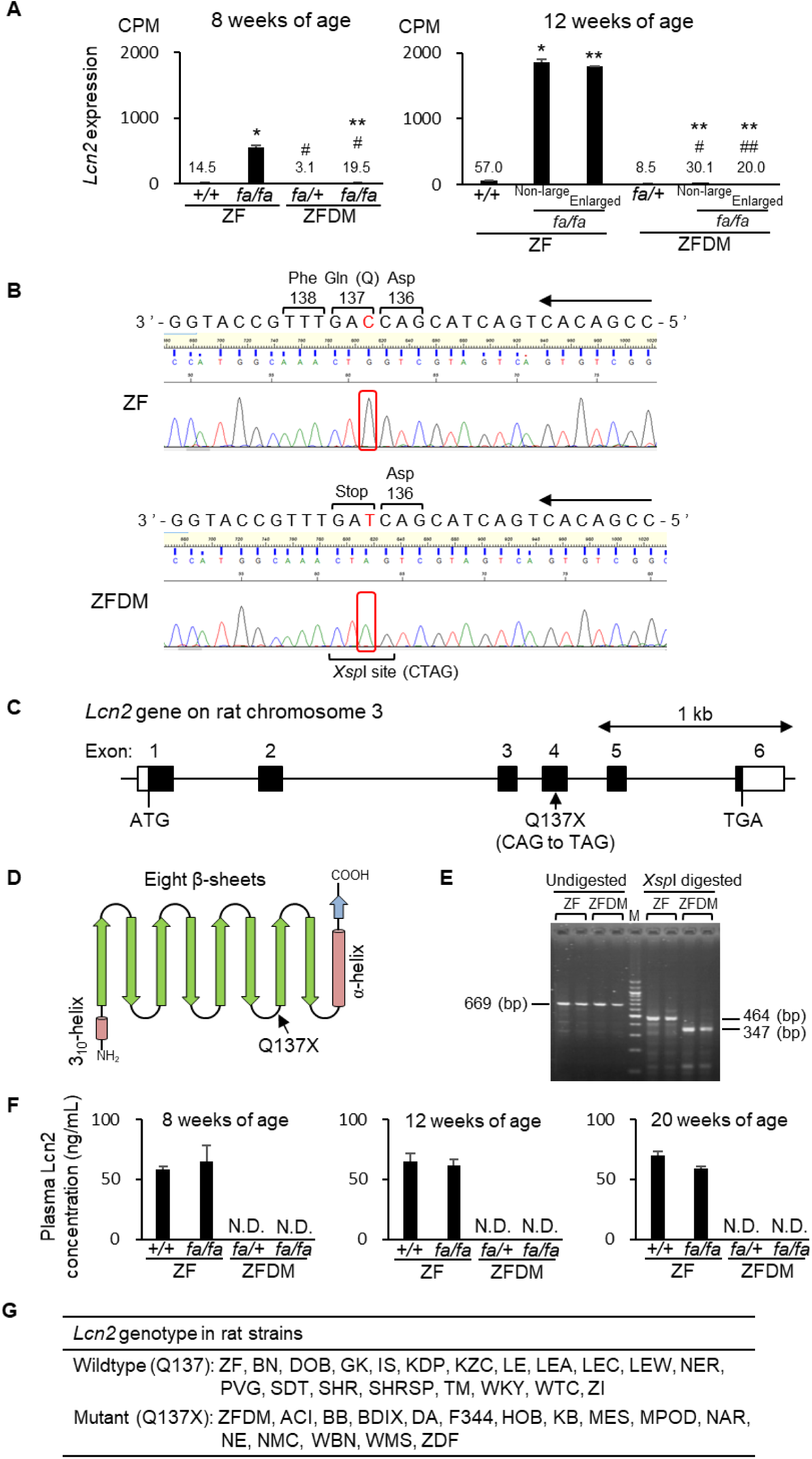
Characterization of the Q137X nonsense mutation in the *Lcn2* gene in ZFDM rats. *(A)* Expression levels of *Lcn* in ZF and ZFDM rats at 8 and 12 weeks of age. The data are CPM values derived from the RNA-sequencing analysis and expressed as means ± SEM (n=3 each). Tukey-Kramer method was used for evaluation of statistical significance: \**P* < 0.05, \*\**P* < 0.01 (vs. +/+ or *fa/+* of each strain); #*P* < 0.05, ##*P* < 0.01 (vs. the corresponding group of ZF rats). *(B)* Electropherogram of the sequencing data containing the Q137X nonsense mutation in the *Lcn2* gene. The mutation produces an *X*spI restriction site. (*C*) Schematic diagram of the exon-intron structure of *Lcn2* gene in rat chromosome 3. (*D*) Schematic diagram of the secondary structure of Lcn2 protein. (*E*) Electrophoretic gel image of the genotyping by PCR-RFLP analysis. *Xsp*I digestion of the 669 bp PCR fragment produces 464 bp and 347 bp fragments in ZF and ZFDM rats, respectively. M, 100 bp DNA ladder marker. (*F*) Plasma Lcn2 levels in ZF and ZFDM rats at 8, 12, and 20 weeks of age. The data are expressed as means ± SEM (n=3 each). Lcn2 protein was not detected (N.D.) in ZFDM rats. (*G*) *Lcn2* genotypes in rat strains. See Supplemental Table S13 for details.

Sanger sequencing confirmed the stop-gained variant in *Lcn2* and verified that ZFDM rat has a nonsense mutation at the 137th glutamine codon (CAG to TAG), c.409C>T; p.Q137X (Fig. 2B). *Lcn2* consists of 6 exons spanning ∼3.3 kb genomic region on rat chromosome 3 and the Q137X mutation is located on exon 4 (Fig. 2C). Lcn2 protein belongs to the lipocalin family, and the protein structure is characterized by a single polypeptide chain that forms a barrel-like structure composed of eight β-sheets, which create a central cavity (Fig. 2D). The mutation disrupts the protein after the 6th β-sheets, deleting 7th and 8th β-sheets and C-terminal α-helix domain. The mutant and wildtype alleles were clearly distinguished by PCR-RFLP analysis (Fig. 2E). Since Lcn2 is known to be secreted into the blood, plasma Lcn2 levels were examined in ZF and ZFDM rats. Plasma Lcn2 proteins were detected in both +/+ and *fa/fa* of the ZF rat with no significant changes with age. In contrast, Lcn2 proteins were hardly detected in plasma of both *fa*/+ and *fa/fa* of the ZFDM rat (Fig. 2F).

To clarify the frequency of the mutation in rat species, we searched for the mutation among 159 inbred and two outbred rat strains. Interestingly, the mutant allele was detected in 32 inbred and one outbred strains including well-known inbred rat strains such as BB, DA, and F344 (Fig. 2G; Supplemental Table S13).

### Evaluation of the relevance of the mutation with T2D by genome editing technique

To evaluate directly that the *Lcn2* mutation is responsible for the development of T2D in ZFDM rats, we generated *Lcn2* knock-in rats to correct the stop-gained variant in *Lcn2* gene in ZFDM rats using CRISPR/Cas9-mediated genome editing.

Among ∼30 founder (G0) animals, there were two animals (G0-28 and -29) harboring an edited allele in which the nonsense mutation was replaced with a wild-type nucleotide (Fig. 3A). Successful correction of the Q137X mutation was confirmed by the fact that animals in both G0-28- and -29-derived lines heterozygous for the edited allele showed the expression of the Lcn2 protein in the plasma (Fig. 3B). We then produced *fa/fa* animals homozygous or heterozygous for the edited allele and compared phenotypes with animals homozygous for the mutant allele (Fig. 3C). There was no significant difference in body weights among animals with the three genotypes. Blood glucose levels also showed no significant difference among the three genotypes. Accordingly, there was no significant difference in cumulative incidence of diabetes among them. Although Lcn2 has been reported to be related with plasma lipid levels (Wallenius et al. 2021), there was no clear difference in plasma lipid parameters among these animals (Supplemental Fig. S1).

**Figure 3.**
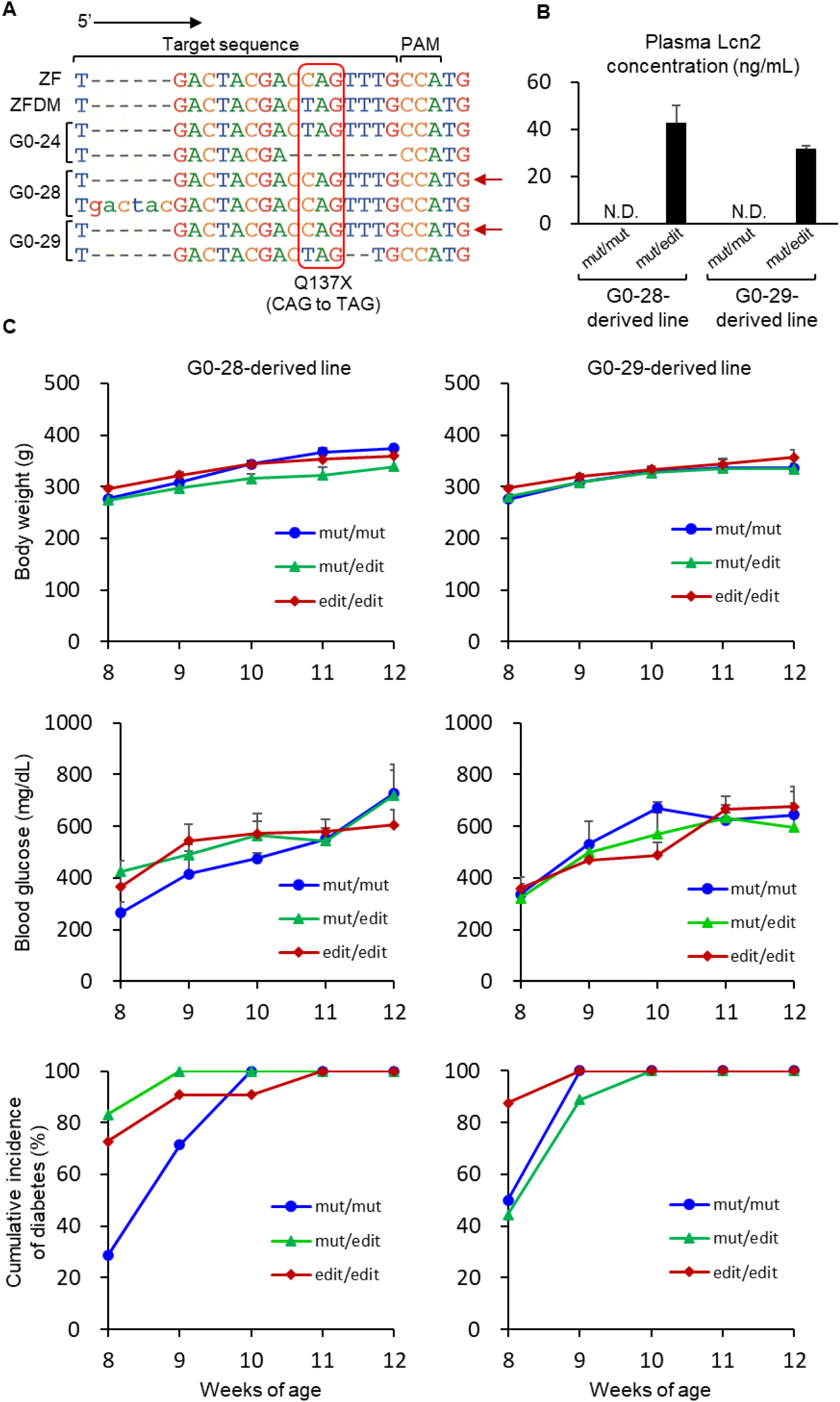
Generation and characterization of *Lcn2* knock-in ZFDM rats. (*A*) Sequence alignment for the target sequence of genome editing. Red arrows indicate the correctly edited alleles. (*B*) Plasma Lcn2 levels in progenies in G0-28- and -29-derived lines: mut, the mutant allele; edit, the correctly edited allele. The data are expressed as means ± SEM (n=5 each, except for mut/edit in G0-29-derived line: n=4). Lcn2 protein was not detected (N.D.) in animals homozygous for the mutant allele. (*C*) Body weights, blood glucose levels, and cumulative incidence of diabetes in G0-28- and -29-derived lines: mut, the mutant allele; edit, the correctly edited allele. The data are expressed as means ± SEM (n=6-12).

### Genetic linkage analysis of the mutation with T2D

To further confirm the relevance of the mutation with T2D, we also performed a genetic linkage analysis by using genetic cross between ZF and ZFDM rats. Since none of male F1 animals homozygous for *fa/fa* developed diabetes, we produced backcross progenies by crossing between female ZFDM and male F1 (Fig. 4A). Among 100 male backcross progenies (*fa/fa*), 57 animals developed diabetes till 60 weeks of age, suggesting that a recessively acting autosomal allele in ZFDM rats is involved in the development of T2D in the cross (Fig. 4B). We determined the *Lcn2* genotypes in each backcross progeny and performed chi-square test between the genotype and diabetic phenotype (Fig. 4C). It has been revealed that there was no significant relationship of the *Lcn2* genotype with the onset of diabetes.

**Figure 4.**
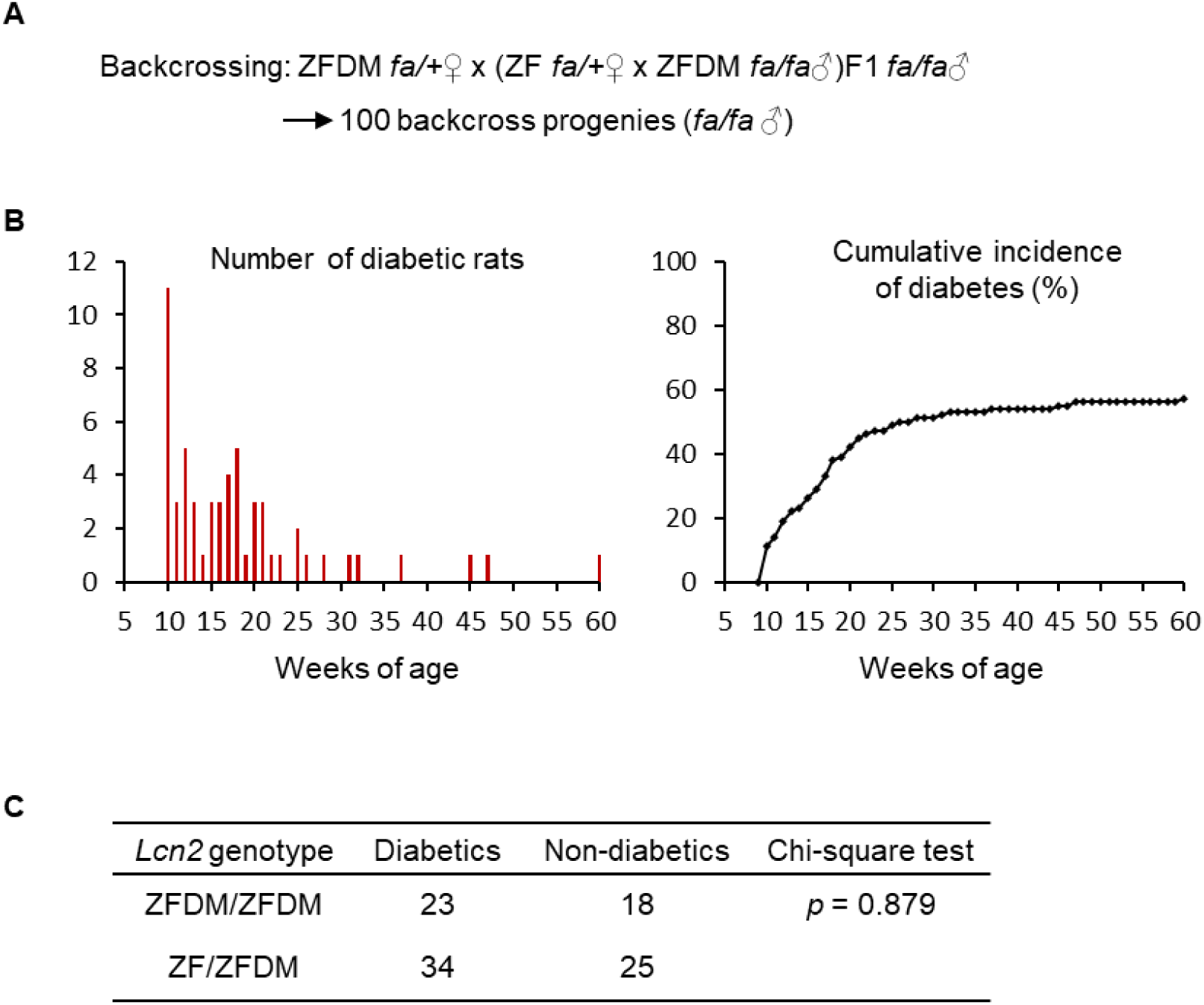
Genetic linkage analysis of the Q137X mutation with the onset of T2D. (*A*) Overview of the method producing backcross progenies between ZF and ZFDM rats. (*B*) Number of diabetic rats (left) and cumulative incidence of diabetes (right) in the backcross progenies till 60 weeks of age. (*C*) Chi-square test for the association between the *Lcn2* genotype and diabetic phenotype in the backcross progenies.

## Discussion

In the present study, by using ZFDM rat as a model of obese T2D, we found that 1) 46 genes exhibit significant expression differences between ZF and ZFDM islets irrespective of obesity and glucose intolerance states; 2) among these common DEGs, the ZFDM rat has a nonsense mutation in *Lcn2* gene, the mutation is distributed widely in rat species; 3) the *Lcn2* mutation is, however, not involved in the development of T2D.

At first, we performed comparative transcriptome analysis of the pancreatic islets between ZF and ZFDM rats at 8 and 12 weeks of age.The common DEGs of the comparison [ZF +/+ vs. ZFDM *fa*/+] and [ZF *fa/fa* vs. ZFDM *fa/fa*] at each age group were associated with “extracellular matrix”: *Fgb, Fgg, Hapln4, Lgals3, Mmp13, Mmp19,* and *P3h2* showed lower expression, while *Adamts16, Ccn4, Col7a1, Col9a1, Cspg4, Cthrc1, Fmod, Olfml2a, Ptx3, S100a4, S100a6, Serpinf1, Spon2, Srpx2, Tgfb1, Tnn,* and *Wnt5a* showed higher expression in ZFDM rats as compared with ZF rats.

Matrix metalloproteinases (MMPs) degrade collagenous extracellular matrix, which are associated with to tissue remodeling and fibrosis. In Zucker diabetic fatty (ZDF) rats, as compared with lean control rats, gene expressions of *Mmp2*, *-12*, and *-14* in diabetic *fa/fa* rats were increased with the onset of islet dysfunction and diabetes (Zhou et al. 2005). In ZFDM rats, we found that most of *Mmps* exhibit higher expressions in *fa/fa* rats as compared with +/+ or *fa/+* rats (Supplemental Fig. S2A). In addition, the expressions of most of *Mmps* are higher in ZFDM rats as compared with those in ZF rats, while the expressions of *Mmp13* and *Mmp19* shows opposite profiles (Supplemental Fig. S2B), indicating that these *Mmps* might have some roles in suppressing the development of islet dysfunction and diabetes.

Gene expression profiles of collagens (Supplemental Fig. S3A), the main component of extracellular matrix, show a similar pattern with those of *Mmps*, indicating that *fa/fa* animals have more extracellular matrix than +/+ or *fa/+* animals, the degree is higher in ZFDM rats as compared with that in ZF rats. The expressions of *Col7a1* and *Col9a1* are significantly higher in ZFDM rats as compared with those in ZF rats (Supplemental Fig. S3B). Since these collagens consist minor collagen components, the roles in the islet dysfunction and diabetes remain to be elucidated. However, the common DEGs for all the comparisons may correspond to primary gene expression difference between strains irrespective of obesity and subsequent glucose intolerance conditions, which could represent primary genetic difference.

Secondly, we therefore searched for functional variants in genes expressed in ZF and ZFDM islets. Among total of 5 stop-gained variants, only *Lcn2* was found to be included in the common 46 DEGs and ZFDM has a Q137X nonsense mutation, serving *Lcn2* as a strong candidate for T2D. The mutation deletes significant C-terminal domains of the protein, which may lead to a loss-of-function of the protein. In addition, the gene expression of the mutant *Lcn2* was strongly reduced due to the nonsense-mediated mRNA decay, resulting in no detectable protein in the plasma. These findings indicate that the Q137X nonsense mutation causes deficiency of the Lcn2 protein.

To our surprise, the mutation is revealed to be distributed widely in inbred rat species including BB, a model of type 1 diabetes (T1D); DA, a model of collagen-induced arthritis (CIA), adjuvant-induced arthritis (AIA), and experimental autoimmune encephalomyelitis (EAE); F344, a well-used control strain; NAR, a model of analbuminemia; WBN, a model of nonobese T2D; ZDF, a model of obese T2D. Especially, the fact that F344 strains have the mutation needs to be noticed since F344 strains frequently serve as background strains for genome editing experiments and as standard strains for preclinical drug safety assessment. Regarding models of diabetes, T1D model KDP rat and nonobese T2D model SDT rat have wildtype alleles, suggesting no apparent association of the mutation with both T1D and T2D.

Finally, we evaluated the relevance of the mutation with T2D by both genome editing and genetic linkage analysis. Our analyses revealed that the mutation is not involved in the development of T2D in ZFDM rats. In addition to adipose tissue, liver, and immune cells that primarily produce Lcn2, various tissues and cells, including pancreatic β-cells, also produce Lcn2 under inflammation or stress conditions (Kim et al. 2023). Lcn2 is thought to have a beneficial role in the regulation of various aspects of energy metabolism: protection from diet-induced obesity and insulin resistance (Guo et al. 2010), high-fat diet-induced adipose tissue remodeling (Guo et al. 2013), and brown fat activation (Zhang et al. 2014). Serum LCN2 levels positively correlates with energy expenditure and fatty acid oxidation in normal weight but not obese women (Paton et al. 2013). In contrast, other reports showed that serum Lcn2 levels are associated with obesity and insulin resistance in humans and mice (Yan et al. 2007; Wang et al. 2007; Rashad et al. 2017).

Mosialou et al. (2017) reported Lcn2 as a bone-derived hormone with metabolic regulatory effects: osteoblast-derived Lcn2 maintains glucose homeostasis by inducing insulin secretion, improves glucose tolerance and insulin sensitivity, and inhibits food intake. In addition, Mosialou et al. (2020) also reported that Lcn2 counteracts metabolic dysregulation in obesity and diabetes, suggesting a distinct beneficial effect of Lcn2 on β-cell function and adaptive β-cell proliferation during toxicity or onset of obesity. They further proposed a model of compensatory homeostatic role of Lcn2, in which Lcn2 counteracts insulin resistance progression, prevents obesity, and suppresses diabetes, while once this mechanism is overwhelmed, obesity increases and diabetes develops. Taken together, our findings in the ZFDM rat, an extreme obese and severe T2D model, suggest that the compensatory homeostatic mechanism is overwhelmed and Lcn2, therefore, could not exert any beneficial effects on β-cell function and proliferation during the development of obesity and T2D. The role of Lcn2 need to be examined in other rat models of mild obesity or diabetes.

In conclusion, we here find a novel nonsense mutation in the *Lcn2* gene in a rat model of obese T2D, the mutation is distributed widely in rat species. Although we could not show the relevance of the *Lcn2* mutation with the development of T2D, several rat strains would serve as *Lcn2* deficient models, contributing to unravel normal and pathophysiological roles of Lcn2 in a wide variety of phenotypes.

## Methods

### Ethics approval and consent to participate

All animal experiments were approved by the Committee on Animal Experimentation of Kobe University and Kyoto University and carried out in accordance with the Guidelines for Animal Experimentation at Kobe University and Kyoto University.

### Animals

For isolation of the pancreatic islets and genetic linkage analysis, male ZF rats (Slc:Zucker-*Lepr^fa/fa^* and -*Lepr^+/+^*) were purchased from Japan SLC, Inc. and male ZFDM rats (Hos:ZFDM-*Lepr^fa/fa^* and -*Lepr^fa/+^*) were provided by Hoshino Laboratory Animals, Inc. All animals were maintained under specific pathogen-free conditions with a 12 h light-dark cycle and were provided with a commercial diet CE-2 (CLEA Japan, Inc.) at the Animal Facility of Kobe Biotechnology Research and Human Resource Development Center of Kobe University. At the end of the experiments, animals were sacrificed by overdose of anesthesia with pentobarbital sodium (2018 or before).

For genome editing experiment, female and male SD rats (Slc:SD) were purchased from Japan SLC, Inc. and female ZFDM rats (Hos:ZFDM-*Lepr^fa/+^*) and male ZFDM rats (Hos:ZFDM-*Lepr^fa/fa^* and -*Lepr^fa/+^*) were provided by Hoshino Laboratory Animals, Inc. All animals were maintained under specific pathogen-free conditions with a 14 h-light and 10h-dark cycle and were provided with a commercial diet F-2 (Oriental Yeast Co., LTD.) at the Institute of Laboratory Animals, Graduate School of Medicine, Kyoto University. At the end of the experiments, animals were sacrificed by carbon dioxide inhalation (2019 or later).

### Isolation of the pancreatic islets

Pancreatic islets were isolated by the collagenase digestion and Ficoll gradient method (Okeda et al. 1979; Carter et al. 2009). Isolated pancreatic islets were cultured for 3 days in RPMI1640 (Sigma-Aldrich) before experiments.

### RNA sequencing and data analysis

Total RNA was extracted from the pancreatic islets using RNeasy Micro kit (Qiagen). RNA sequencing (125 bp paired-end) was performed on 1 μg each of total RNA, using an Illumina HiSeq 2500 system by Eurofins Genomics. Sequence reads were cleaned using trimmomatic (ver. 0.39) (Bolger et al. 2014), the quality were checked using FastQC (version 0.11.9) (Andrews S 2010), and then aligned to the rat genome (mRatBN7.2) using STAR (version 2.7.10) (Dobin et al. 2013). Data were transformed into BAM format using Samtools (version 1.15.1) (Danecek et al. 2021), and raw read counts were calculated using featureCounts (version 2.0.3) (Liao et al. 2014). The following analyses were performed using R (version 4.1.3) (https://www.r-project.org/). Filtering low expression genes, TMM normalization, and extraction of DEGs were done using edgeR (version 3.40.1) (Robinson et al. 2010; Chen et al. 2016). DEGs were extracted using quasi-likelihood F-test of edgeR with threshold of FDR < 0.01 and fold change > 2. Information of DEGs was obtained using biomaRt (version 2.54.0) (Durinck et al. 2005; Durinck et al. 2009). GO and pathway analyses were performed using DAVID (Huang et al. 2009; Sherman et al. 2022). The RNA sequencing data have been deposited in DDBJ Sequence Read Archive (DRA) with the accession number DRA007109 (Hayami et al 2020), DRA007371 (Hayami et al 2020), and DRA012479.

### Variant search using RNA sequencing data

SNPs between ZF and ZFDM rats were detected from BAM files of RNA sequencing data of the pancreatic islets at 8 weeks of age using GATK (version 4.2.6.1) (McKenna et al. 2010) with the GATK Best Practice (DePristo et al. 2011; Van der Auwera and O’Connor 2020). Resulting Variant Calling Format (VCF) files were summarized using VCFtools (version 0.1.16) (Danecek et al. 2011). Variants were annotated using VEP (version 108) (McLaren et al. 2016). SNPs were defined as those fixed for distinct homozygous state between ZF and ZFDM rats.

### Mutation screening of the lipocalin 2 gene

Genome DNA was extracted from tail tip samples using the Wizard SV Genomic DNA Purification System (Promega). All amino acids coding regions in lipocalin 2 gene of ZF and ZFDM rats were sequenced by the Sanger method. A PCR-RFLP system was developed for genotyping of the Q137X nonsense mutation found in ZFDM rats: PCR product (669 bp) amplified by using a primer set (FW, 5’-aaccctgggtatgacctgaa-3’; RV, 5’-ctggggcctggattattgta-3’) is digested by restriction enzyme *Xsp*I, resulting in 464 bp, 114 bp, and 91 bp fragments in ZF rats (wildtype allele) while the 464 bp fragment is further divided into 347 bp and 117 bp fragments in ZFDM rats (mutant allele). The PCR-RFLP system was also applied for genotyping of the mutation among rat species. The genome DNA of 157 inbred rat strains (Supplemental Table S13) were provided by the National Bioresource Project-Rat (NBRP-Rat), Kyoto University (Kyoto, Japan). The genome DNA of the SDT/Jcl rat has been obtained previously (Fuse 2008). The ZDF-*Lepr^fa^*/CrlCrlj rat was purchased from the Jackson Laboratory Japan, Inc. and genome DNA was extracted.

### CRISPR/Cas9-mediated genome editing in ZFDM rats

*Lcn2* knock-in ZFDM rats, in which the Q137X nonsense mutation was replaced with a wild-type nucleotide, were generated by CRISPR/Cas9-mediated genome editing as described previously with some modification (Yoshimi et al. 2016). Briefly, female ZFDM *fa*/+ rats were superovulated by intraperitoneally injection with 15 IU of pregnant mare serum gonadotropin (PMSG: NIPPON ZENYAKU KOGYO Co., Ltd.) and 15 IU of human chorionic gonadotropin (ASKA Pharmaceuticals Co., Ltd.), and then the female rats were mated with male ZFDM (*fa*/+ or *fa/fa*) rats. The next day, pronuclear-stage embryos were collected from superovulated rats and cultured in a modified Krebs-Ringer bicarbonate (m-KRB) culture medium before microinjection. The *Lcn2* target sequence (5’-TGACTACGACTAGTTTGCCA-3’; Fig. 3A) was designed using CRISPOR (http://crispor.gi.ucsc.edu). The recombinant Cas9 protein and crRNA and tracrRNA were purchased from Integrated DNA Technologies (Coralville, IA, USA). The chemically synthesized single-strand oligo-DNA (ssODN; 5’-AAGTGGCCGACACTGACTACGACCAGTTTGCCATGGTATTTTTCCAGAAGACCTCTGAAA-3’, Exigen) was used for replacing the nonsense mutation with the wild-type allele on the *Lcn2* locus. The recombinant Cas9 protein (50 ng/mL), chemically synthesized crRNA (25 ng/mL) and tracrRNA (25 ng/mL), and the ssODN (100 ng/μL) were co-injected into the cytoplasm of pronuclear stage embryos. The injected embryos were cultured in m-KRB culture medium overnight. The two-cell embryos were transferred into the oviduct of pseudopregnant SD rats anesthetized using mixture of 0.15 mg/kg medetomidine, 2.0 mg/kg midazolam and 2.5 mg/kg butorphanol during operation. After the surgery, 0.15 mg/kg atipamezole (NIPPON ZENYAKU KOGYO Co., Ltd.) and 5 mg/kg enrofloxacin (Kyoritsu Seiyaku) were administered. Analgesics (0.01 mg/kg buprenorphine; Otsuka Pharmaceutical Co. Ltd.) were administered subcutaneously twice on the day of surgery and the following day.

To confirm the genome edited alleles, the sanger sequencing analysis was performed on the target locus. The PCR products were obtained from tail tip genome DNAs by using the primer set (FW, 5’-aaccctgggtatgacctgaa-3’; RV, 5’-ctggggcctggattattgta-3’), and sequenced with each of the primers. Two female founder rats (G0-28 and -29) heterozygous for the genome-edited allele were obtained and were mated with male ZFDM rats to produce offsprings. The inheritance of the genome-edited allele was confirmed in the offsprings. Both lines were maintained for several generations to produce *fa/fa* animals homozygous or heterozygous for the genome-edited allele and those homozygous for the original mutant allele. Finally, the genome-edited allele was fixed in the homozygous state. The G0-28- and -29-derived genome-edited ZFDM rats, ZFDM-*Lcn2^em1Nyo^*and -*Lcn2^em2Nyo^*, were deposited to the NBRP-Rat under deposition No.0972 and No.0973, respectively.

### Phenotyping

The rats were checked for body weights and nonfasting blood glucose levels by a portable glucose meter (ANTSENSE Duo, HORIBA, Ltd.). Diabetes was defined as blood glucose levels equal to or higher than 300 mg/dL for three consecutive weeks under ad libitum dietary conditions. The week of the diabetes onset was defined as the first week in which blood glucose levels were equal to or higher than 300 mg/dL.

### Genotyping of the *fatty* mutation in the leptin receptor gene

A PCR-RFLP system was used for genotyping the *fatty* mutation. PCR product (596 bp) amplified by using a primer set (FW, 5’-aagccatctcatttgctggt-3’; RV, 5’-ggcaggcagatctctcaatc-3’) is digested by restriction enzyme *Msp*I (CCGG). The 596 bp fragment (wildtype allele) is further divided into 328 bp and 268 bp fragments in the *fatty* allele.

### Plasma Lcn2 levels

Plasma Lcn2 levels were measured using ELISA kits (BioPorto Diagnostics A/S).

### Plasma lipid parameters

Plasma lipid parameters were measured using the medium-sized biochemistry automatic analyzer 7180 (Hitachi, Ltd.).

### Genetic analysis

Female ZF *fa*/+ rats were crossed with male ZFDM *fa/fa* rats to produce F1 animals. Then, female ZFDM *fa*/+ rats were crossed with the male *fa/fa* F1 rats to produce backcross progenies. Male *fa/fa* backcross progenies were checked for nonfasting blood glucose levels until 60 weeks of age. Genotyping of the Q137X nonsense mutation was performed by the PCR-RFLP system as described above.

### Statistical analysis

Data are expressed as mean ± SEM. Differences among the groups were analyzed with Tukey-Kramer method as indicated in the figure legends. Association of the *Lcn2* genotype with diabetes was analyzed with chi-square test. *P* < 0.05 was regarded as statistically significant. Statistical analysis was performed using R (version 4.4.1).

## Data access

All raw RNA sequencing data have been deposited in the DDBJ Sequence Read Archive (DRA; https://www.ddbj.nig.ac.jp/dra/index-e.html) under accession number DRA007109, DRA007371, and DRA012479.

## Acknowledgements

We thank Hoshino Laboratory Animals, Inc. for providing ZFDM rats; Shihomi Hidaka, Ayako Kawabata, Takuro Yamaguchi, and Chihiro Seki for their excellent technical assistance. We also thank the National BioResource Project-Rat (http://www.anim.med.kyoto-u.ac.jp/NBR/) for providing genome DNA of 157 inbred rat strains. This study was supported by JSPS KAKENHI Grant Number JP18H02364 and JP21H02390 (to N.Y.), and partially supported by grants from the National Center for Global Health and Medicine Grant Number 29-1001 and 23A1013 (to T.O.).

## Author contributions

N.Y. conceived the study. N.Y., N.A., T.H., S.S., R.U., and Y.T. performed experiments. N.Y., K.N., Y.Y., S.Y., and T.O. performed genome editing. N.Y., T.H., M.Y., and M.S. performed statistical and bioinformatic analyses. N.Y., M.S., and T.O. supervised the project. N.Y. and N.A. wrote the manuscript. N.Y. edited the manuscript, with input from all authors.

## Supplemental materials

**Supplemental Table S1.** List of 1,046 DEGs for ZF +/+ vs. ZFDM *fa/+* at 8 weeks of age.

**Supplemental Table S2.** List of 359 DEGs for ZF *fa/fa* vs. ZFDM *fa/fa* at 8 weeks of age.

**Supplemental Table S3.** List of 1,355 DEGs for ZF +/+ vs. ZFDM *fa/+* at 12 weeks of age.

**Supplemental Table S4.** List of 778 DEGs for non-large islets of ZF *fa/fa* vs. ZFDM *fa/fa* at 12 weeks of age.

**Supplemental Table S5.** List of 766 DEGs for enlarged islets of ZF *fa/fa* vs. ZFDM *fa/fa* at 12 weeks of age.

**Supplemental Table S6.** List of the common 127 DEGs at 8 weeks of age.

**Supplemental Table S7.** List of GO terms enriched for the 127 DEGs at 8 weeks of age.

**Supplemental Table S8.** List of the common 166 DEGs at 12 weeks of age.

**Supplemental Table S9.** List of GO terms enriched for the 166 DEGs at 12 weeks of age.

**Supplemental Table S10.** List of the common 46 DEGs for all the comparisons.

**Supplemental Table S11.** List of GO terms enriched for the common 46 DEGs for all the comparisons.

**Supplemental Table S12.** List of variants found on RNA-seq data of the islets of ZF and ZFDM rats at 8 weeks of age.

**Supplemental Table S13.** *Lcn2* genotypes among rat strains.

**Supplemental Figures**

## Description of supplemental figures

**Supplemental Fig S1.** Plasma lipid parameters in G0-28- and -29-derived lines.

**Supplemental Fig S2.** Gene expression profiles of matrix metalloproteinases (MMPs) in ZF and ZFDM rats at 12 weeks of age.

**Supplemental Fig S3.** Gene expression profiles of collagens in ZF and ZFDM rats at 12 weeks of age.

