## Supplemental Figures for "Comparative transcriptome and variant analyses of the pancreatic islets of a rat model of obese type 2 diabetes identifies a frequently distributed nonsense mutation in the lipocalin 2 gene"

**Description of supplemental figures**

**Supplemental Fig S1. Plasma lipid parameters in G0-28- and -29-derived lines.**

**Supplemental Fig S2. Gene expression profiles of matrix metalloproteinases (MMPs) in ZF and ZFDM rats at 12 weeks of age.**

**Supplemental Fig S3. Gene expression profiles of collagens in ZF and ZFDM rats at 12 weeks of age.**

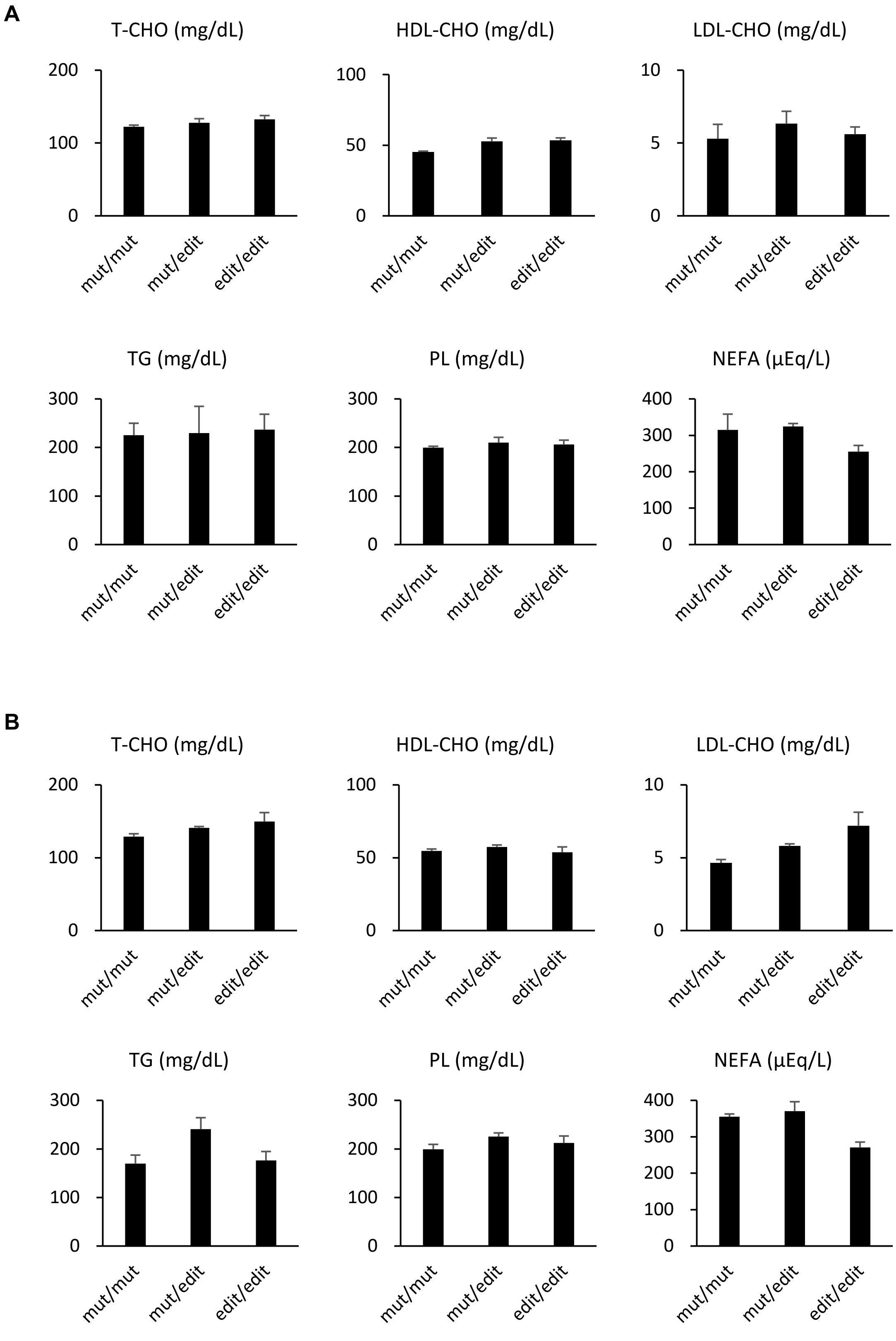


**Supplemental Figure S1. Plasma lipid parameters in G0-28- and -29-derived lines.** (*A*) Plasma lipid parameters in G0-28-derived lines. (*B*) Plasma lipid parameters in G0-29-derived lines. Data are expressed as means ± SEM (n=4-5 each). mut, the mutant allele; edit, the correctly edited allele. T-CHO, total cholesterol; HDL-CHO, HDL cholesterol; LDL-CHO, LDL cholesterol; TG, triglyceride; PL, phospholipid; NEFA, non esterified fatty acid.

**A**

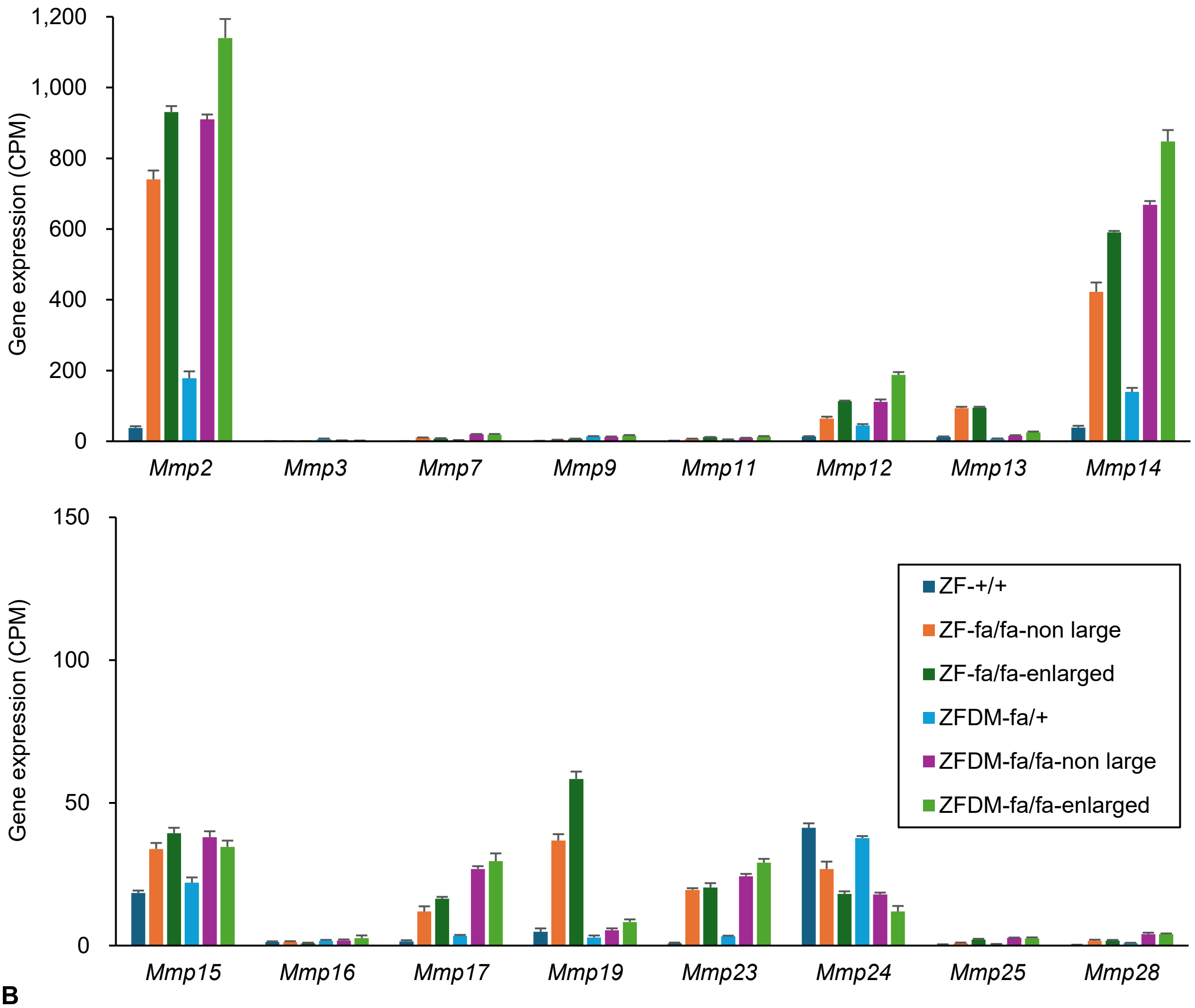


**B**

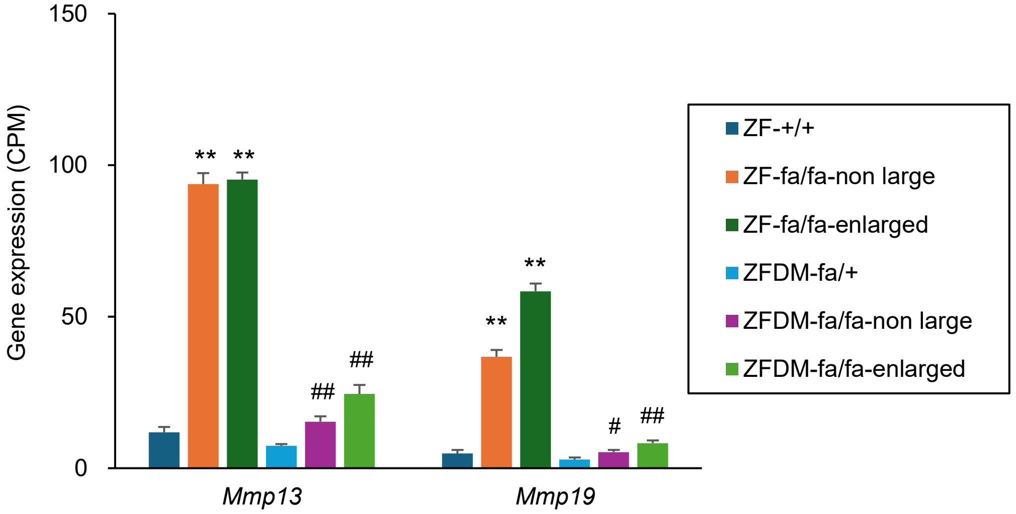


**Supplemental Figure S2. Gene expression profiles of matrix metalloproteinases (MMPs) in ZF and ZFDM rats at 12 weeks of age.** (*A*) Expression levels of *Mmps* in ZF and ZFDM rats at 12 weeks of age. The data are CPM values derived from the RNA-sequencing analysis and expressed as means ± SEM (n=3 each). (*B*) Expression levels of *Mmp13* and *Mmp19* in ZF and ZFDM rats at 12 weeks of age. The data are CPM values derived from the RNA-sequencing analysis and expressed as means ± SEM (n=3 each). Tukey-Kramer method was used for evaluation of statistical significance: ***P* < 0.01 (vs. +/+ or *fa*/+ of each strain); #*P* < 0.05, ##*P* < 0.01 (vs. the corresponding group of ZF rats).

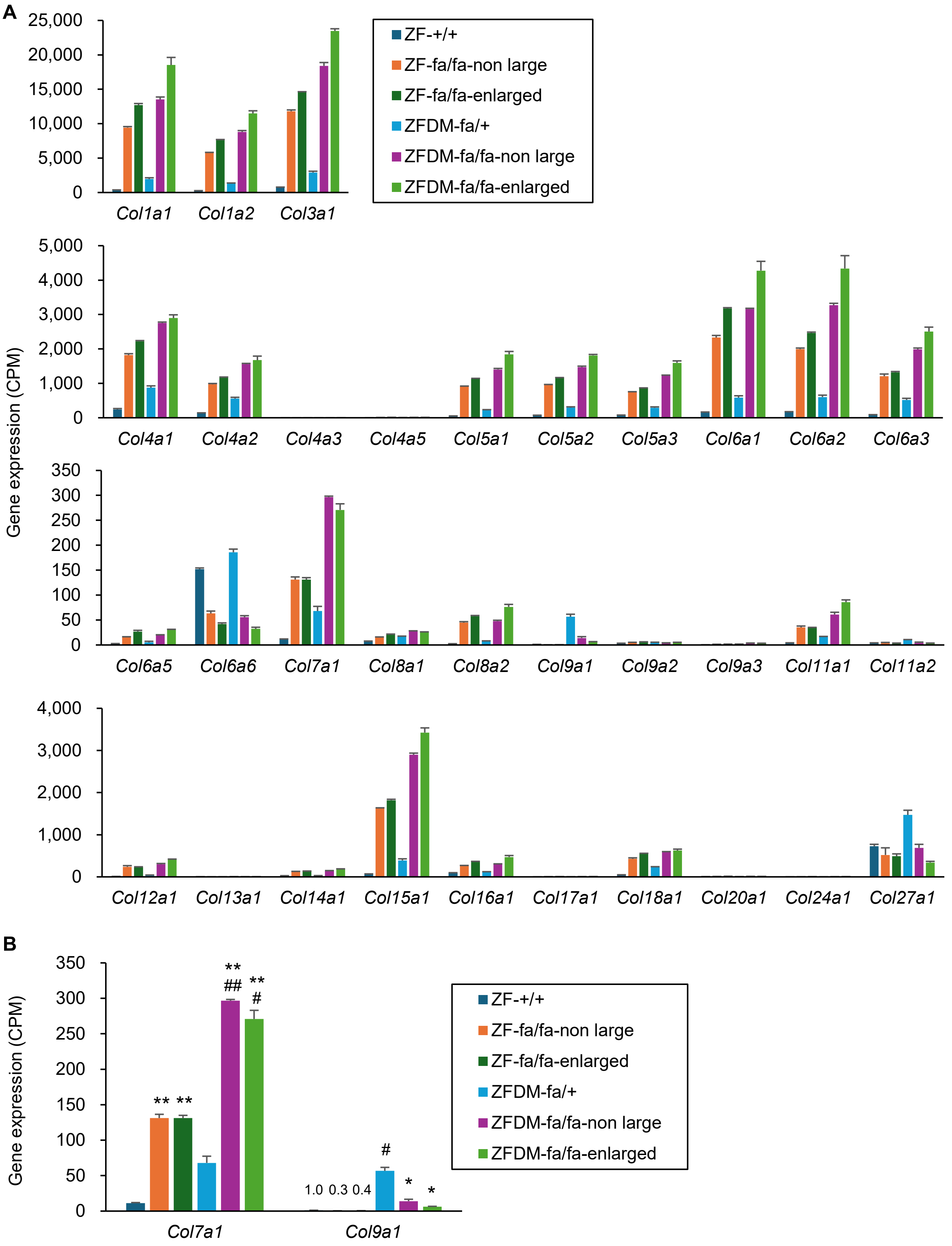


**Supplemental Figure S3. Gene expression profiles of collagens in ZF and ZFDM rats at 12 weeks of age.** (*A*) Expression levels of collagens in ZF and ZFDM rats at 12 weeks of age. The data are CPM values derived from the RNA-sequencing analysis and expressed as means ± SEM (n=3 each). (*B*) Expression levels of *Col7a1* and *Col9a1* in ZF and ZFDM rats at 12 weeks of age. The data are CPM values derived from the RNA-sequencing analysis and expressed as means ± SEM (n=3 each). Tukey-Kramer method was used for evaluation of statistical significance: *$P < 0.05$, **$P < 0.01$ (vs. +/+ or *fa*/+ of each strain); #$P < 0.05$, ##$P < 0.01$ (vs. the corresponding group of ZF rats).
